# Deep-learning based denoising and reconstruction of super-resolution structured illumination microscopy images

**DOI:** 10.1101/2020.10.27.352633

**Authors:** Zafran Hussain Shah, Marcel Müller, Tung-Cheng Wang, Philip Maurice Scheidig, Axel Schneider, Mark Schüttpelz, Thomas Huser, Wolfram Schenck

## Abstract

Super-resolution structured illumination microscopy (SR-SIM) provides an up to two-fold enhanced spatial resolution of fluorescently labeled samples. The reconstruction of high quality SR-SIM images critically depends on patterned illumination with high modulation contrast. Noisy raw image data, e.g. as a result of low excitation power or low exposure times, result in reconstruction artifacts. Here, we demonstrate deep-learning based SR-SIM image denoising that results in high quality reconstructed images. A residual encoding-decoding convolution neural network (RED-Net) was used to successfully denoise computationally reconstructed noisy SR-SIM images. We also demonstrate the entirely deep-learning based denoising and reconstruction of raw SIM images into high-resolution SR-SIM images. Both image reconstruction methods prove to be very robust against image reconstruction artifacts and generalize very well over various noise levels. The combination of computational reconstruction and subsequent denoising via RED-Net shows very robust performance during inference after training even if the microscope settings change.

## 1 Introduction

Fluorescence microscopy remains one of the most powerful tools for imaging cell biology samples because of its ability to specifically label molecular structures and to visualize them in different fluorescence color channels. It also offers exceptionally high sensitivity and can visualize molecular processes even below the optical diffraction limit. Several methods have been developed during the last two decades [1] that enable imaging of fluorescently labeled samples down to the nanoscale. Super-resolution structured illumination microscopy (SR-SIM) is a particularly compelling method, because it works with the majority of samples and fluorophores commonly used in cell biology without imposing specific requirements to sample preparation [2,3]. Therefore, it can even be applied to living samples [4–12]. SR-SIM in its most common form utilizes a series of sinusoidal illumination patterns with a pattern periodicity at or near the diffraction limit. This patterned excitation light is phase-shifted laterally and rotated to different discrete angles to acquire a series of raw images, which are then passed on to an image reconstruction algorithm to obtain the final super-resolved SR-SIM image [13]. Several implementations of reconstruction algorithms that operate in frequency space were developed and a number of open-access tools are now available that aim to enhance the speed, spatial resolution and minimize reconstruction artifacts in the final reconstructed image, i.e. fairSIM [14], OpenSIM [15], SIMToolbox [16], and CC-SIM [17]. The most common property among all these image reconstruction algorithms is that they require a series of high-quality raw images in order to be able to reconstruct a high-quality super-resolution image. However, reconstruction in frequency space fails to reliably reconstruct SR-SIM images if the signal-to-noise ratio (SNR) is too low, e.g. because the laser excitation power level was too low, the sample exposure time was chosen too short, or the sample has already undergone irreversible photobleaching [18].

Recently, several developments with the aim to reduce reconstruction artifacts from SR-SIM images have been undertaken. Huang et al. used a fully analytical approach to reduce noise and minimize reconstruction artifacts using a Hessian matrix approach [10]. Hoffman and Betzig used the reconstruction of SIM images in lower pixel count tiles and subsequent merger to reduce reconstruction artifacts [19]. Jin et al., on the other hand, used deep neural networks to reconstruct the cropped regions of SR-SIM images [20]. Christensen et al. used a deep learning architecture for the reconstruction of synthetic SR-SIM images [21] with subsequent testing on real microscope images. Ling et al. relied on a special type of convolutional neural network, a CycleGAN, for the same purpose [22]. Weigert et al. used deep learning algorithms to enhance isotropic resolution and signal-to-noise ratio of fluorescence microscopy images in general [23].

In our work, we first suggest an end-to-end deep learning architecture and workflow which is related to the work by Jin et al. [20] but uses a different network architecture and works directly on full-size microscope images (also in contrast to [21] where synthetic data is used for training). Furthermore, as completely novel workflow, we combine the classical computational SIM reconstruction with subsequent denoising via a deep network. Therefore, the focus of our work is to demonstrate and compare (1) the direct reconstruction of SIM raw images with a low signal-to-noise ratio into high-quality SR-SIM images and (2) the denoising and artifact reduction of low SNR SR-SIM images.

The first approach is named super-resolution residual encoder-decoder structured illumination microscopy (SR-REDSIM), the second approach is named residual encoder-decoder fairSIM (RED-fairSIM). Both are used to reconstruct super-resolved images from raw noisy SIM input images, but they differ in their use of deep learning: SR-REDSIM is an end-to-end deep learning model, where the full SIM reconstruction process is performed by the deep learning network. RED-fairSIM, on the other hand, is the combination of the fairSIM image reconstruction package, which performs image reconstruction using the widely used frequency-domain algorithms, and then subsequently uses a deep learning network for artifact reduction and denoising. We found that both of these methods are robust in their ability to significantly improve the quality of reconstructed SR-SIM images. We also demonstrate that both these methods are robust in reconstructing SR-SIM images from noisy raw SIM images with different noise levels. Similarly, we also showed, that the RED-fairSIM generalizes better even if the microscope settings are changed after training.

## 2 Results

Deep learning methods rely on training data, which in our case consist of noisy raw SIM images as input and ideally noise and artifact free, super-resolved SIM reconstructions as output. Thus, we first need to generate such a data set that is large enough to effectively train the network, but also captures all aspects of the SIM imaging process (sample behaviour, instrument imperfections, data processing artifacts) well enough.

In principle, the data acquisition process of a SIM microscope can be simulated. In this case, the expected output represents the ground truth data on which the simulation is based upon, and which is known without SNR or resolution limits. Also, the amount of available training data would only be limited by processing time, as the generation would be fully automated and not rely on access to a microscope system.

However, we have decided against this *in silico* approach. While the basic effects of structured illumination, Poisson-distributed noise and even basic optical imperfections are rather easy to simulate, modelling the response of a full structured illumination microscope correctly is very complex. Additionally, such a simulation would likely have to be adjusted to reflect the properties of a specific SR-SIM instrument in order to capture changes, e.g. when switching to a different manufacturer or even a specific installation of an SR-SIM microscope. The same argument holds for the fluorescent samples themselves. While some simulations providing perfect ground-truth data exist, e.g. for single molecule localization microscopy [24,25], they, again, do not capture all of the variability found in real-world samples.

The option chosen for data generation for the work presented here is to use the real microscope and standard biological samples. This approach naturally captures all aspects and imperfections of the samples and of the specific instrument in question [23], but also poses constraints. As data collection requires both instrument time and manual sample preparation and handling, the amount of training data is naturally limited. Also, there is no *perfect* ground-truth available. To acquire the high-quality reference images presented to the networks as desired output, we thus adjusted the instrument to provide high SNR raw frames and processed those with the classical, frequency-domain-based image reconstruction algorithm. While these images are low in noise and reconstruction artifacts, they are never completely devoid of them. To acquire noisy, low-SNR images as input, the samples were then photo-bleached by continued exposure and acquisition of raw SIM images, which naturally reduces fluorescent light output over time and results in a series of images with steadily decreasing SNR (see 4.2 for details).

### 2.1 SR-REDSIM: SR-SIM image denoising and reconstruction using the super-resolution REDSIM method

In the first deep-learning based SR-SIM image reconstruction method, named SR-REDSIM, both the tasks of reconstruction and denoising of noisy raw SIM images are performed by a single deep learning model. This model is a modified version of the “Residual Encoder-Decoder Network” (RED-Net). RED-Net is an encoding-decoding framework with symmetric convolutional-deconvolutional layers along with skip-layer connections. It was previously used to accomplish different image restoration tasks such as image denoising, image super-resolution, and image inpainting [26]. The original RED-Net architecture is only composed of encoding-decoding blocks with the size of the network input being the same as the size of the network output. Therefore, super-resolution with this architecture has to rely on explicit image pre-upsampling [26]. In contrast, our modified RED-Net architecture contains an additional upsampling block after the encoding-decoding blocks. This upsampling block inside our model has the advantage that the input images are first denoised in their lower-dimensional space which reduces training time and effort.

Most of the super-resolution architectures such as the enhanced deep super-resolution network (EDSR) [27] or the residual channel attention network (RCAN) [28] are very deep and require a significant amount of training data. In comparison, our architecture is comparably light-weight.

The complete pipeline of this method is shown in Fig. 1 and the architecture of SR-REDSIM (see supplementary Fig. 1) is explained in more detail in section 4.3. During the training process, we used all 15 raw noisy SIM images (3 angles with 5 phases each) of size 512 × 512 pixels (i.e., stack dimensions were 15 × 512 × 512 (frames, width, height)) as input along with the reconstructed super-resolved SIM image of size 1024 × 1024 pixels as output. The output was generated by the fairSIM software from raw SIM images recorded with the best signal-to-noise ratio, while the input images were taken from noise level 4 (for an explanation of the noise levels see section 4.2). The trained network was afterwards tested on unseen test data from noise level 4. The super-resolution images obtained during this test are depicted in column 2 of Fig. 2 whereas column 1 and 5 show the results of “noisy fairSIM” (reconstructed by fairSIM from noisy raw SIM images; noise level 4) and “reference fairSIM” (reconstructed by fairSIM from raw SIM images with the highest SNR). The comparison of these images and of specific regions of interest (ROIs) between fairSIM in Fig. 2 (column 1, all rows) and SR-REDSIM (column 2, all rows) clearly shows that the noise is completely removed by SR-REDSIM. However, in the reconstruction by SR-REDSIM fine cell structures are partly suppressed as compared to the reference output (compare column 2, row 2, ROI 1 in Fig. 2 with column 5, row 2, ROI 1). Anyhow, in rows 3/4 and 5/6 of Fig. 2 the structure of the cell is well denoised and reconstructed by SR-REDSIM. Moreover, the evaluation of the SR-REDSIM method on the basis of peak single to noise ratio (PSNR) and structural similarity index measurement (SSIM) [29] values in Table 1 shows a significant improvement compared to fairSIM.

**Figure 1:**
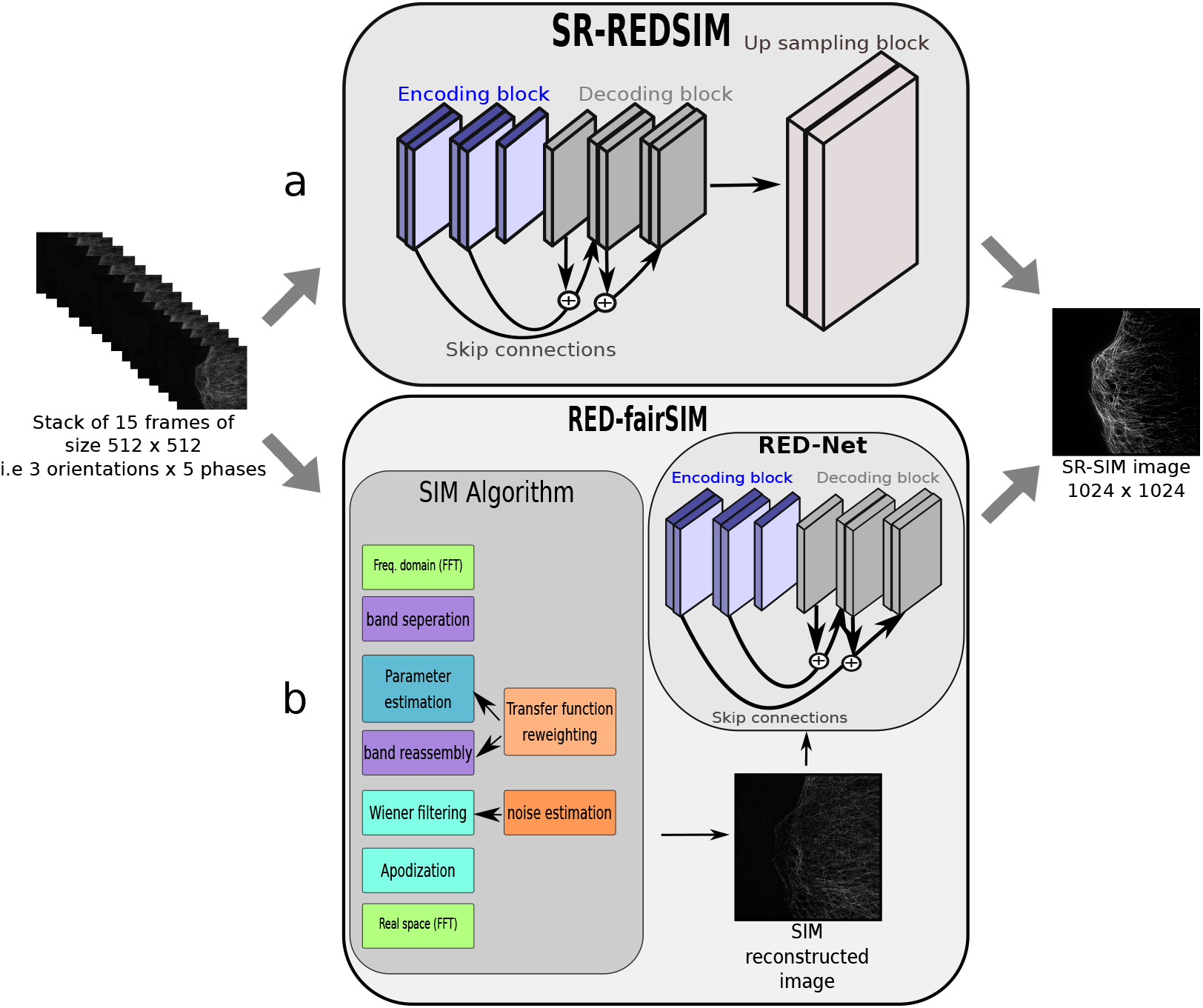
Schematics of the deep learning CNN architecture of the two different SR-SIM image denoising (RED-fairSIM) and image denoising and reconstruction (SR-REDSIM) methods. In both approaches a stack of 15 raw (noisy) SIM images (3 angles with 5 phases each) is used as input. The output is the reconstructed SR-SIM image. **a** SR-REDSIM is composed of three main blocks. The encoding block contains mainly the convolutional layers whereas the decoding block consists of deconvolutional layers and the up-sampling block of deconvolutional up-sampling layers. **b** In the RED-fairSIM method, fairSIM is first used to computationally reconstruct noisy SR-SIM images which are then further propagated into the RED-Net for denoising. The architecture of RED-Net is composed of the encoder and the decoder block.

**Figure 2:**
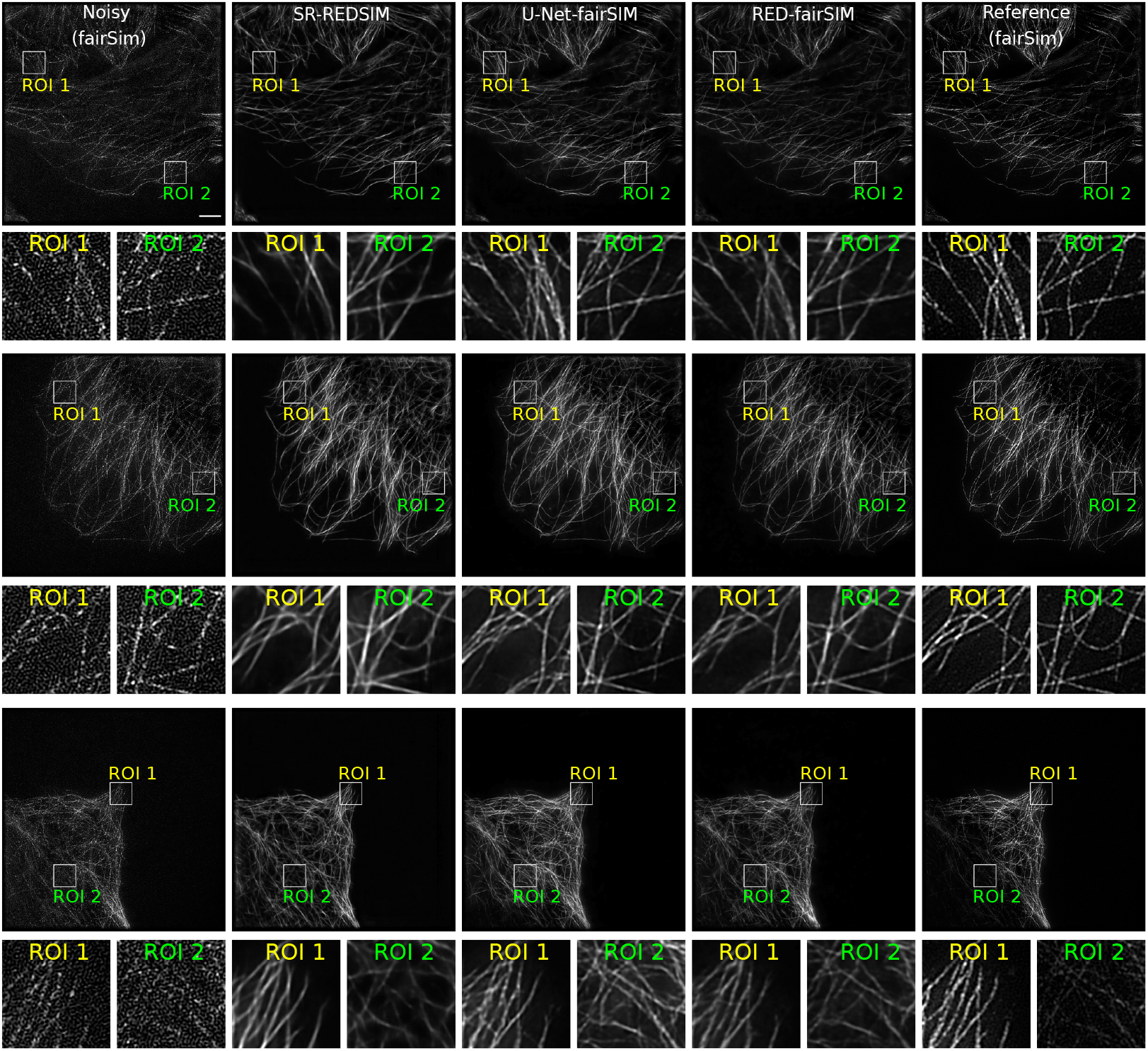
Super-resolution SIM (SR-SIM) images of three different test samples (U2OS osteosarcoma cells, tubulin cytoskeleton labeled with anti-tubulin-Alexa488) at high-level noise (time stamps 175 – 199; noise level 4). Each column represents a different reconstruction approach: fairSIM (first column), SR-REDSIM (second column), U-Net-fairSIM (third column), and RED-fairSIM (fourth column). The fifth column depicts the reconstructed reference images which were generated by applying fairSIM image reconstruction to high SNR image data at noise level 0 (lowest noise level, i.e. timestamp 0). All reconstructed SR-SIM images have 1024 × 1024 pixels. The first, third, and fifth rows correspond to the full-size SR-SIM images, whereas the second, fourth, and sixth rows depict magnified regions of interest (ROIs) of the white squares (bounding boxes) indicated in the full-size images. The extracted ROIs of size 100 pixels × 100 pixels were upsampled to 300 pixels × 300 pixels using bicubic interpolation for illustration purposes. Scale bar: 4 μm.

**Table 1:** Mean PSNR and SSIM values of all methods (computed with the fairSIM output at noise level 0 as reference on the test data)

|  | PSNR and SSIM values at different noise levels |  |  |  |  |  |  |  |
| --- | --- | --- | --- | --- | --- | --- | --- | --- |
|  | noise level=1 |  | noise level=2 |  | noise level=3 |  | noise level=4 |  |
|  | PSNR | SSIM | PSNR | SSIM | PSNR | SSIM | PSNR | SSIM |
| fairSIM | 28.84 | 0.69 | 25.83 | 0.45 | 24.12 | 0.32 | 23.61 | 0.29 |
| SR-REDSIM | 26.73 | 0.69 | 26.62 | 0.64 | 26.66 | 0.64 | 26.62 | 0.69 |
| U-Net-fairSIM | 27.44 | 0.75 | 27.23 | 0.69 | 26.85 | 0.65 | 26.80 | 0.68 |
| RED-fairSIM | 28.75 | 0.80 | 28.67 | 0.75 | 28.18 | 0.70 | 27.97 | 0.71 |

### 2.2 RED-fairSIM: SR-SIM reconstruction of noisy input data by using the combination of fairSIM and RED-Net

fairSIM is one of the well known open-source reconstruction algorithm implementations and is widely used for super-resolution tasks among the other tools in SIM microscopy. However, it cannot reconstruct a clean high-quality super-resolution image from noisy raw SIM images. In the RED-fairSIM method, fairSIM is first used to reconstruct the SIM samples of noisy raw SIM images and then RED-Net is used to denoise the output of fairSIM and to generate in this way high-quality super-resolution images. During the reconstruction process, again the stack of 15 noisy images (3 angles, 5 phases) of size 512 × 512 is propagated into the fairSIM reconstruction algorithm which then generates a single noisy reconstructed image of size 1024 × 1024 pixels. The noise and artifacts found in these images do not follow a typical distribution (e.g. Poisson or Gaussian), but have a distinct form which comes from the frequency-based reconstruction algorithm. This single noisy reconstructed images is further passed into the RED-Net architecture to achieve the final result that can be seen in Fig. 2 (column 4). The complete pipeline of the RED-fairSIM method can be seen in Fig. 1 and the architecture of RED-Net is shown in supplementary Fig. 2.

The parameters which were used to generate SIM reconstructed samples from the raw SIM images are explained in section 4.4.1. RED-Net was trained in a supervised way where the input-output pairs contain the noisy and reference reconstructed images. The noisy reconstructed images are selected from noise level 4. For the output image we used the term reference instead of ground truth because the reconstruction by fairSIM shows artifacts even for rather clean raw images.

The performance of this method on the unseen test samples is the best among our experiments with respect to PSNR and SSIM values (see table 1) as well as visually. The regions of interest (ROIs) in Fig. 2 show clearly that the output images generated by RED-fairSIM are of high quality with fine details and smooth lines. They are superior compared to “noisy fairSIM”, “reference fairSIM”, and SR-REDSIM. Even the artifacts introduced by fairSIM in the reference images are completely removed by RED-fairSIM.

Furthermore, preliminary tests of RED-fairSIM and SR-REDSIM concerning their ability to generalize to different SIM imaging conditions were carried out (see supplementary Fig. 3). As before, U2OS cells were stained for microtubuli, but a dark-red dye with illumination shifted to 642nm was used, which subsequently also shifts the spatial frequencies of the illumination pattern. The RED-fairSIM approach is able to denoise these images and remove SIM reconstruction artifacts, while the SR-REDSIM approach creates heavy ghosting artifacts. This is unsurprising, as in the case of SR-REDSIM, all specific properties of the SIM pattern (spatial frequencies, orientation, phases) are learned by the network. In the RED-fairSIM approach, parameters specific to the SIM pattern are absorbed by the classic, frequency-domain-based reconstruction, and only reconstruction artifacts are carried into the network. Those artifacts might still depend on the SIM imaging parameters, so further cross-checks should be carried out. As an initial result, RED-fairSIM seems to generalize well to different SIM pattern settings.

**U-Net-fairSIM** U-Net is also a popular learning architecture that is extensively used in the domain of image restoration such as image denoising and super-resolution [23,30,31]. In the pipeline proposed for RED-fairSIM, we replaced the RED-Net by U-Net and analyzed the resulting super-resolution images. The architecture of U-Net is shown in supplementary Fig. 4. Henceforth, we named this approach as U-Net-fairSIM. Figure 2 (column 3) shows that U-Net-fairSIM also produces better results as compared to the noisy, reference and SR-REDSIM images. However, it does not surpass RED-fairSIM. Similarly, the U-Net-fairSIM approach outperforms all other counterparts except for RED-fairSIM concerning the PSNR and SSIM values in table 1. Comparing RED-fairSIM and U-Net-fairSIM directly as in Fig. 3, the cell structures reconstructed by RED-fairSIM are smoother. Furthermore, they are more faithful when taking the reference as “gold standard” into account. For these reasons, we have focused the presentation in this paper on RED-fairSIM.

**Figure 3:**
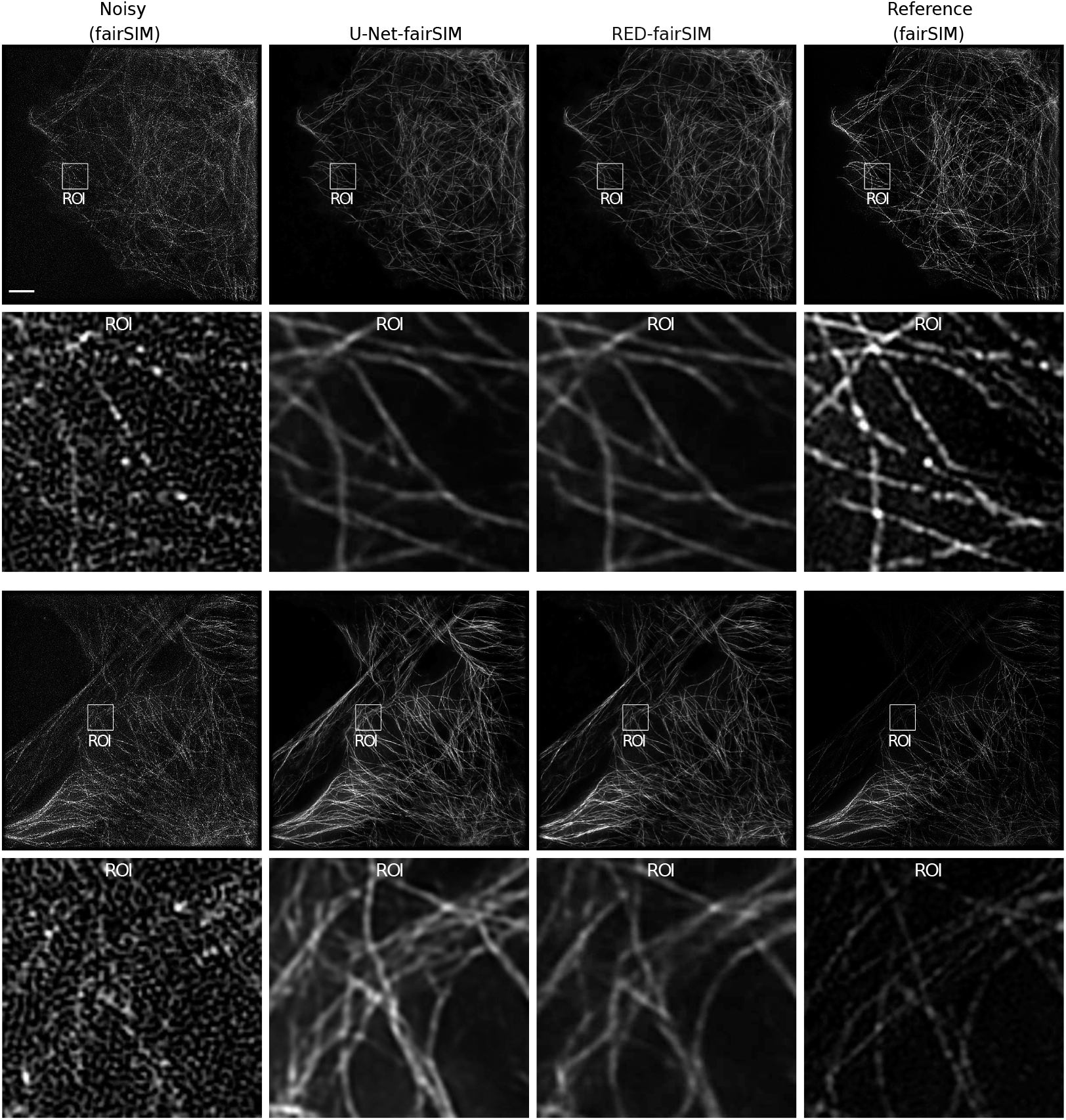
Reconstruction of SR-SIM images of two different test samples with the fairSIM, U-Net-fairSIM and RED-fairSIM methods. Each column represents the results of the corresponding method. The first and third row show the resulting SR-SIM images whereas the second and fourth row contain the extracted enlarged regions of interest (ROIs) from the full-size images in the row directly above. The cell structures reconstructed by RED-fairSIM are smoother compared to U-Net-fairSIM and fairSIM (reference). Furthermore, they are more faithful than U-Net-fairSIM when taking the reference as “gold standard” into account. Scale bar: 4μm.

During all of the reported experiments, we did not carry out any preprocessing on the input or output images such as downsampling or cropping (see section 4.2). Nonetheless, we tested if image augmentation yields any significant improvement of the results. For this purpose, each image was rotated by an angle of 180° and added in this form to the training set. This increased the amount of training data from 2025 to 4050 images. The results with and without image augmentation are reported in supplementary table 1. Because image augmentation did not provide a noticeable advantage overall and did not change the performance-wise ordering of the proposed methods, but on the other hand doubles the training time and increases the preprocessing effort, we decided to focus in this paper on the results without image augmentation.

### 2.3 Alternative Approaches

**preRED-fairSIM** Similarly, we also tried to generate high-quality super-resolution SR-SIM images by another method named as preRED-fairSIM. The pipeline of preRED-fairSIM is shown in the supplementary section (see supplementary Fig. 5). However, preRED-fairSIM failed to deliver usable results in the end. In preRED-fairSIM, each noisy SIM image from a different phase and orientation is denoised separately and then the whole stack of all 15 denoised images is propagated into the fairSIM algorithm to reconstruct a final super-resolution image. In the preRED-fairSIM approach we trained a 30-layer RED model for one selected phase and orientation and then performed transfer learning [32] (which implies in our scenario no re-training and no changes in the network weights) and fine tuning [33] (which implies adaptation of a subset of the network weights) to other phases and orientations. The results of transfer learning and fine-tuning are quite promising in terms of the achieved SSIM and PSNR values (see supplementary table. 2). The empirical results of this approach on the image level are shown in Fig. 4 (with fine-tuning applied). They also prove that the raw SIM images are well denoised in the first step of this method. However, fairSIM fails to reconstruct super-resolution images of sufficient quality from the denoised raw images. The resulting reconstructed images contain some additional new artifacts. These artifacts can likely be traced to higher harmonics introduced by the RED model in the preRED denoising step, which become very clear in the Fourier power spectrum of the denoised images (see fig 4), and then appear similarly as artifacts in the Fourier spectrum of the fully reconstructed image. This is to be expected, as the fairSIM method works in the frequency domain, and highly relies on the precise phases, orientations and harmonics of the SIM pattern, which the denoising step obviously breaks.

**Figure 4:**
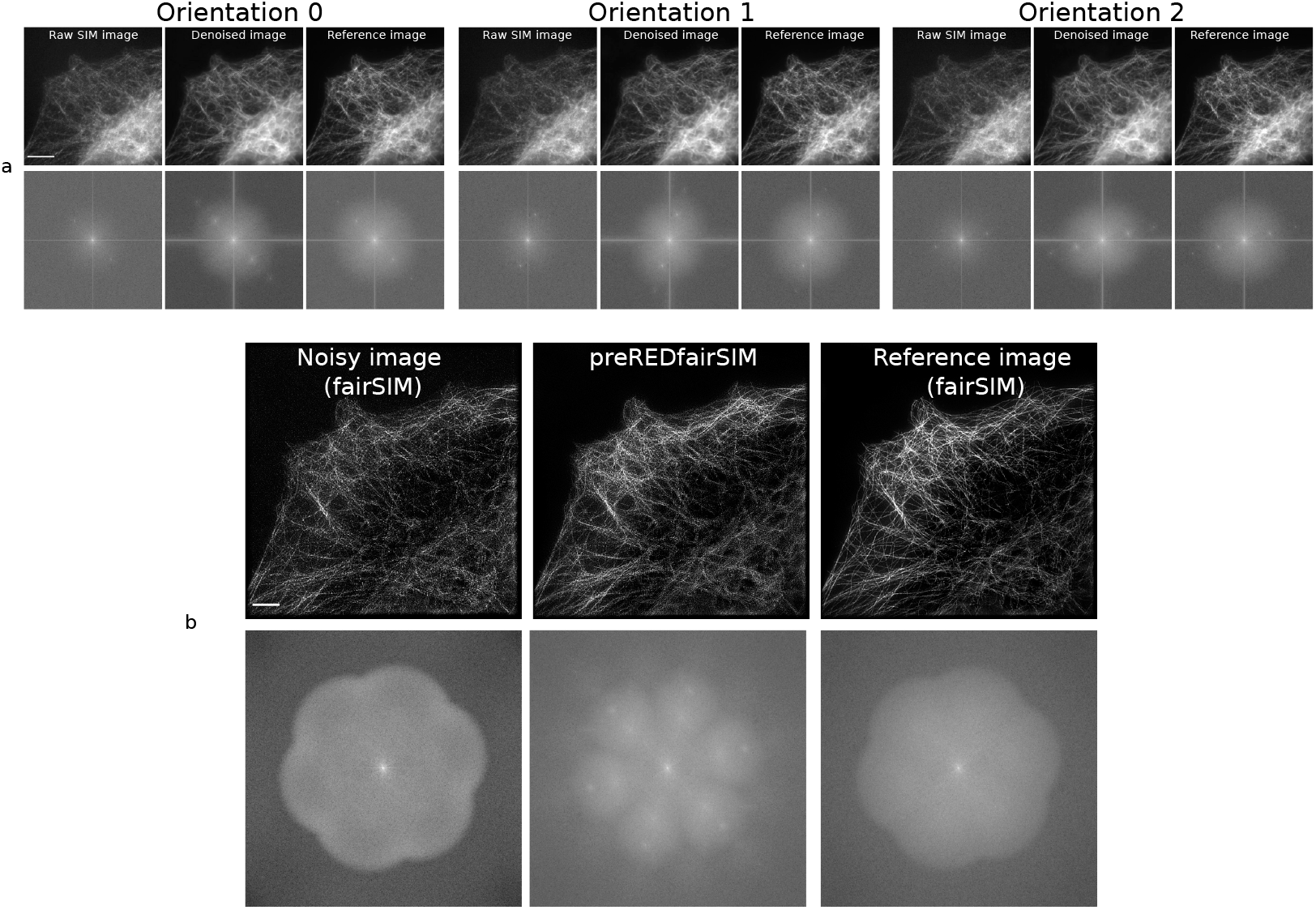
preRED-fairSIM results. **a** shows three blocks of images where each block consists of 6 images. The first block depicts the images from phase 0 and orientation 0, the second block from phase 1 and orientation 1, and the third block from phase 2 and orientation 2. The left image in the first row of each block represents a noisy raw SIM image from noise level 4. The second image in each block is the denoised version whereas the reference image (rightmost in each block) is the ground truth. The Fourier spectra of the images are shown below each image. The dimensions of each image in these blocks are 512 × 512 (width × height). Scale bar: 8 μm. **b** shows three images reconstructed by fairSIM along with their Fourier spectrum directly below. The noisy image (left) is reconstructed using 15 noisy raw SIM images and the parameter fit summary is: resolution improvement (*x* :1.90, *y* :1.90, *z* :1.90), modulation estimation (*x* :0.310, *y* :0.341, *z* :0.332) with assessment as “weak”. Similarly, the preRED-fairSIM image (middle) is generated using 15 denoised SIM images and the the parameter fit summary is: modulation estimation (*x* :0.310, *y* :0.341, *z* :0.332) with assessment “weak”, however, there is no improvement in the resolution. The reference image (right) is reconstructed from the raw SIM images with the highest SNR and the parameter fit summary is: resolution improvement is (*x* :1.90, *y* :1.90, *z* :1.90), modulation estimation (*x* :0.558, *y* :0.580, *z* :0.578) with assessment “usable”. The Fourier spectrum of preRED-fairSIM shows additional artifacts (white spots) that neither exists in the Fourier spectrum of the reference nor the noisy output. Scale bar: 4 μm.

**BM3D** BM3D is a conventional state-of-the-art image denoising method from the field of computer vision [34]. During this work, we also used BM3D to denoise the noisy super-resolution images reconstructed by fairSIM. BM3D is able to remove the noise successfully from reconstructed noisy SIM samples (see supplementary Fig. 6, column 3) but fails to recover lost information.

### 2.4 SIM reconstruction at varying noise levels

The raw SIM data in this study was collected as a time-lapse of a fixed sample undergoing photo-bleaching. Thus, data was collected at different noise levels, which we can be assembled into five noise level groups, with *noise level*=0 representing the lowest and *noise level*=4 representing the highest level of noise found in the data. As we have previously discussed, the models were trained with input from the highest noise level and therefore, we investigated whether these pre-trained models will also be useful for the SIM images of other, lower noise levels. To verify this, we considered only the two best methods from our work: SR-REDSIM and RED-fairSIM are used to evaluate the raw SIM images at different levels. No fine-tuning [33] or transfer learning [32] were performed on these pre-trained models. Figure 5 shows the results of this attempt. In this figure, one specific sample was captured at different noise levels. If we further examine the ROIs of all the super-resolution images, it can be seen that both methods show high-quality super-resolved images at all of the five different noise levels with a slight degradation towards higher noise levels regarding smoothness and clarity of the cell structures. Furthermore, it is again noticeable that the results of RED-fairSIM are overall visually more appealing compared to the other methods. In addition to visual inspection, quantitative results are given in table 1 which contains the mean PSNR and SSIM values of 500 test inputs from each noise level. Here, U-Net-fairSIM is included in the presented comparison. Both RED-fairSIM and U-Net-fairSIM show a gradual decrease in PSNR and SSIM values from noise level 1 (light noise) to noise level 4 (strong noise). SR-REDSIM performs similarly, but with noticeably smaller PSNR and SSIM values. The results of fairSIM without denoising deteriorate quickly when moving to higher noise levels. The most important take-away from table 1 and Fig. 5 are that the networks — although trained for a specific high noise level — generalize well to conditions with a better signal-to-noise ratio.

**Figure 5:**
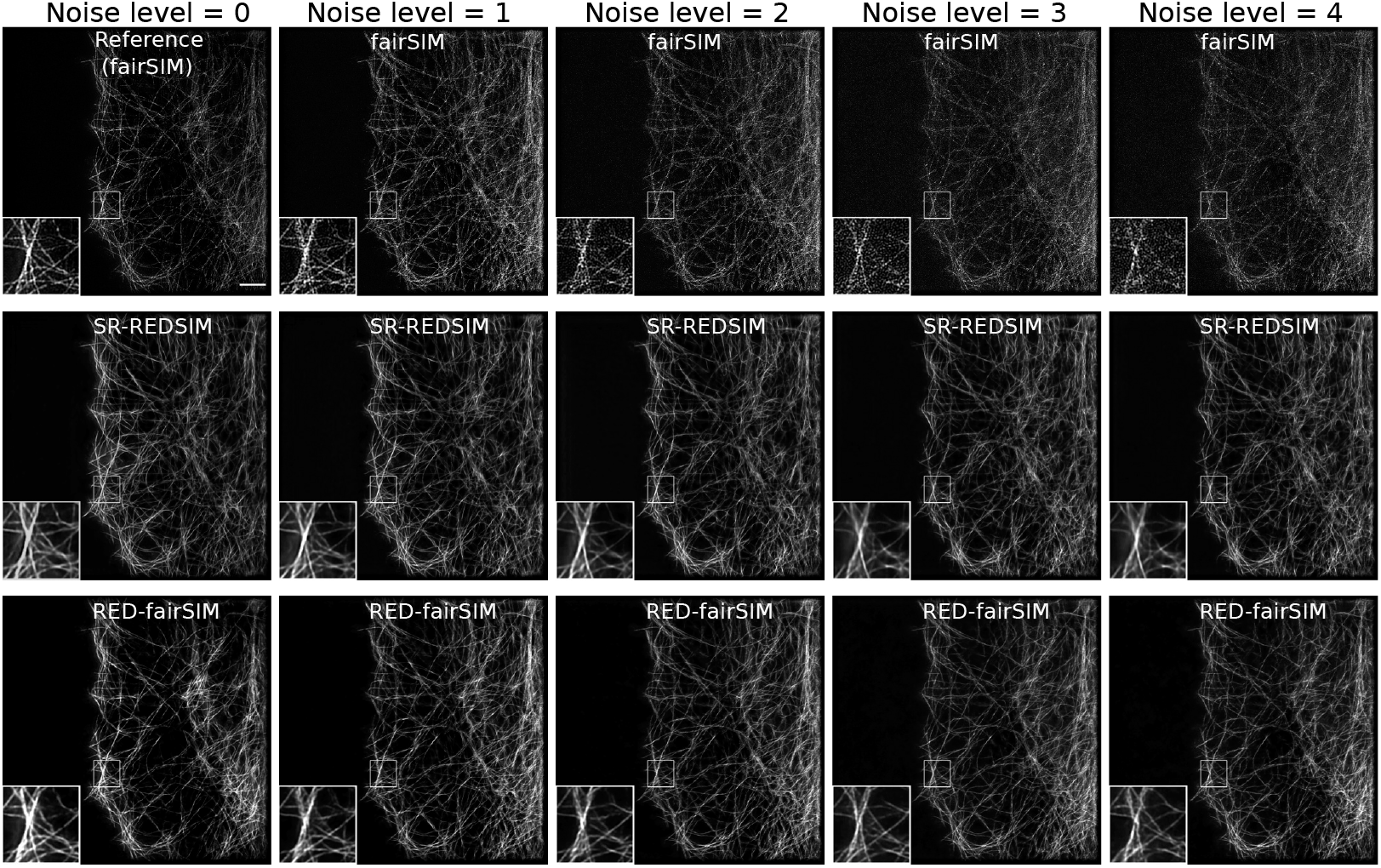
Reconstructed SR-SIM images at different noise levels with the SR-REDSIM and RED-fairSIM methods for a single field of view. Noise level 0 represents the reference image at timestamp 0, noise level 1 comprises the images from timestamps 25 – 50, noise level 2 from timestamps 75 – 100, noise level 3 from timestamps 125 – 150, and noise level 4 from timestamps 175 – 200. Each full image contains an enlarged region of interests (ROI) in the bottom left. The images reconstructed by fairSIM in the first row show a significant degradation in quality as the noise level increases. In contrast, the results produced by SR-REDSIM and RED-fairSIM in the second and third are far less depending on the noise level. Scale bar: 4 μm

The reconstruction of the lower noise level 0 through SR-REDSIM and RED-fairSIM highlights a second use case. At this noise level, the SNR of the raw frames is high enough to provide the reference data sets, which as discussed before are of high quality, but still feature some SIM reconstruction artifacts. Those artifacts are successfully removed by both SR-REDSIM and RED-fairSIM. A reasonable assumption is that, like noise, reconstruction artifacts are random enough in nature, so they are not picked up by the network during training, and thus cannot be reproduced. This effect is well known in applications such as noise2noise [35], where the inability of a neural network to learn random (noisy) data is explicitly used for denoising.

## 3 Discussion

The results of this study provide sufficient evidence that SR-REDSIM and RED-fairSIM can both be employed to denoise and reconstruct high-quality SR-SIM images. We also investigated the robustness of the proposed methods (SR-REDSIM and RED-fairSIM) and showed that high-quality reconstruction of SIM samples is possible irrespective of the noise level in the raw SIM images. The SR-REDSIM and RED-fairSIM methods outperform their counterparts as shown in Fig. 2 and supplementary Fig. 6. Furthermore, these approaches are useful even in the absence of clean groundtruth data as we have shown especially for RED-fairSIM where the reference data used for training contains many reconstruction artifacts. We have also shown in Fig. 5 (column 1) that the proposed methods SR-REDSIM and RED-fairSIM can be used to remove the reconstruction artifacts from the reference image after training, so even if high SNR data can be acquired easily, SR-REDSIM and RED-fairSIM still offer an improvement over the classical reconstruction approaches.

A recent study [22] used cycle-consistent generative adversarial networks (CycleGAN) [36] to reconstruct SR-SIM images by using 3 to 9 clean raw SIM images. A CyleGAN contains two generators and two discriminators with multiple losses that are trained in a competitive process. Therefore, CycleGANs are generally very difficult to train. Furthermore, the authors did not address the challenge of denoising. Christensen et al. [21] trained deep neural networks by using synthetic data instead of real microscope SIM images to reconstruct SR-SIM images. Although the synthetic data used in their studies for training is unrelated to real microscopes, they were successful in generating output comparable to computational tools like fairSIM. However, they did not use real noisy microscope data for their testing of the denoising performance of their networks, and their approach was also not completely successful in the case of (simulated) high-level noise. Jin et al. [20] used multiple concatenated U-Nets to reconstruct SR-SIM images by using 3 to 15 raw SIM images. They trained their models on cropped and resized SIM samples and manually discarded tiles with only background information. These preprocessing steps are time-consuming, and the training of two adjacent U-Net model is also computationally expensive.

Our proposed methods use the raw SIM images in their original size which does not involve any major preprocessing steps. The amount of training data used, about 100 fields of view for training and test data together, is also small enough that specific training, capturing both a given instrument and a specific biological structure of interest, should often be feasible. While SR-REDSIM has similarities to other proposed end-to-end deep learning approaches for SIM [20–22], RED-fairSIM is a completely novel deep learning approach for SIM which is — as our data shows — superior to SR-REDSIM.

While both SR-REDSIM and RED-fairSIM provide high-quality reconstruction, an obvious difference between them is their ability to generalize to different SIM imaging settings. As an initial test, we varied the spatial frequency of the SIM pattern (using 642nm instead of 488nm excitation light), which commonly happens when designing experiments and choosing dyes (see supplementary Fig. 3). We then reconstructed with RED-fairSIM and SR-REDSIM, both trained on the original 488nm data. Here, the RED-fairSIM approach, where the change in spatial frequency of the pattern is absorbed by the classic reconstruction step, still works very well in suppressing noise and SIM artifacts. SR-REDSIM, on the other hand, where the SIM pattern has been learned by the network, created heavy ghosting artifacts. While further validation and cross-testing is needed, this suggests that RED-fairSIM should be able to generalize to different SIM microscopes, excitation wavelength and probably illumination types (3-beam, 2-beam, TIRF-SIM), while SR-REDSIM would require retraining whenever larger changes in these parameters occur.

## 4 Methods

### 4.1 Sample preparation and data acquisition

U2OS cells were cultured in DMEM supplemented with 10% FBS and grown on round coverslips of 170±5 nm thickness (No. 1.5H). Cells were fixed with 4% PFA for 15 min, followed by PBS washes, permeabilization with 0.5% Triton-X100 for 3 min. Another two rounds of PBS washes were done prior to blocking with 3% BSA. For the immunolabeling of microtubuli, cells were stained with anti-tubulin Ab (Invitrogen CatNo. 322500) 1:400 for 2 hr at room temperature, followed by PBS wash and one additional hour of incubation with Alexa 488-conjugated anti-mouse IgG 1:400. Cells were then briefly washed with PBS before Vectashield was applied to embed the coverslip onto the standard slide glass for imaging.

The Delta Vision|OMX (GE Healthcare) was used to acquire 3D-SIM raw images. A total of 101 randomly selected field-of-views were acquired by exposing each field-of-view to the full laser power of the 488 nm excitation laser and the exposure time was set at 20 ms for each of the 15 raw image frames. 200 image repetitions were collected at each position without delays. Taking into account camera readout and pattern switching time, acquiring 15 raw SIM images, making up one timestamp in the raw image stack, takes approximately 375 ms.

### 4.2 Data preprocessing

In our work, we used 101 different cell structures (fields of view). Out of these 81 were selected for the training data and the remaining 20 for the test data. Each cell structure was captured for 200 repetitions. During each repetition, 15 frames were recorded, iterating the phase and orientation of the sinusoidal SIM illumination pattern, yielding an image stack of size 15 × 512 × 512 (frames, width, height). During this time-lapse acquisition, the samples underwent photo-bleaching, which reduces the amount of active fluorescent emitters and thus the amount of emitted photons. Therefore, less and less light is captured and the SNR steadily decreases during the acquisition of such a time series.

The cell structures from the timestamps 175 to 200 are therefore considered as noisy training and test input while the samples from timestamp 0 are considered as clean output images. All 15 clean raw SIM images of the 101 cell structures from timestamp 0 are used to reconstruct high-resolution reference SIM images of size 1024 × 1024 pixels by using fairSIM, which employes a classic frequency-domain-based reconstruction. In this work, the input dimension is 15 × 512 × 512 (frames, width, height) whereas the output dimension is 1024 × 1024 (width, height) pixels. The total of 2525 samples were further divided into training and test data. The traininig data contains 2025 images of the first 81 cell structures whereas the test data is composed of 500 test images which are created from the remaining 20 cell structures.

The only preprocessing step which is involved in our work is the linear scaling of the training and test data to match overall brightness between input and output. In addition, we tested an image augmentation approach to double the amount of training data by rotating each image by an angle of 180°.

#### 4.2.1 Definition of noise levels

The data from each time series over 200 repetitions was subdivided into noise levels. Noise level 0 stands for the highest SNR in our data at timestamp 0. This is our reference data. The image data from timestamps 175 – 200 represents the highest noise level 4, the data from timestamps 125 – 150 represents noise level 3, timestamps 75 – 100 noise level 2 and timestamps 25 – 50 noise level 1. In this study, data from noise level 4 is only used in the training process whereas data from noise levels 1, 2, 3, and 4 is used in the test phase.

### 4.3 Architecture and training of SR-REDSIM

SR-REDSIM is based on a modified version of RED-Net. The complete pipeline of this approach is shown in Fig. 1a whereas the architecture of SR-REDSIM and details about the model parameters are given in supplementary Fig. 1. The SR-REDSIM architecture consists of three blocks which are encoder, decoder, and the upsampling block. SR-REDSIM contains in total 44 convolutional and deconvolutional layers with symmetric skip connections.

The encoder block is composed of 21 convolutional layers whereas the decoder contains 21 deconvolutional layers. The upsampling block consists of 2 deconvolutional layers that perform the upsampling task by adjusting the size of the stride. The SR-REDSIM model provides the best results after training the model for 100 epochs. The SR-REDSIM model is trained only with high-level noise data from timestamps 175 to 200. During the training process, the ADAM optimizer and the L2 loss function, also known as Least Squares error, are used:

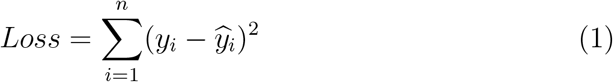

In equation 1, *y_i_* represents the true pixel intensity whereas 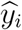 represents the predicted pixel intensity. *n* is the number of pixels (in the image).

### 4.4 Architecture and training of RED-fairSIM

The pipeline of RED-fairSIM is shown in Fig. 1b. In this approach, first fairSIM is used to transform the raw SIM images into a super-resolved output image, by employing a classic, frequency-domain SIM reconstruction algorithm. This output image now contains noise, which due to the frequency-domain algorithm takes a SIM-specific form, and might show other reconstruction artifacts. It is subsequently processed by RED-Net.

#### 4.4.1 fairSIM

The fairSIM reconstruction is performed in three steps, parameter estimation, reconstruction and filtering. Mathematical and algorithmic details are provided in the original publication [14]. A synthetic optical transfer function, with NA = 1.4, λ = 525 nm, *a* = 0.31 (*a* is a compensation parameter, see [14, 37]) is used. 500 counts of background are subtracted per pixel. SIM reconstruction parameters (pattern orientation, global phase, etc.) are automatically determined by fairSIMs standard, iterative cross-correlation approach. Filter parameters are set to a generalized Wiener filter strength of *w* = 0.05, apodization is set at 1.9× the resolution limit with a *bend* of 0.8, and a notch-style filter implemented as *OTF attenuation* with a strength of 0.995 and a FWHM of 1.2 μm^−1^ is in use. For the detailed meaning and influence of these parameters, please refer to the fairSIM source code, its accompanying publication [14] and this general guide to SIM reconstruction parameters [37].

#### 4.4.2 RED-Net Architecture

The architecture of RED-Net as used in RED-fairSIM consists of 15 convolutional and 15 deconvolutional layers along with symmetric skip connections (see supplementary Fig. 2). The output of fairSIM is propagated into the RED-Net to denoise the reconstructed noisy sample as shown in Fig. 2 (column 1). During the training phase, the noisy SR-SIM images of size 1024 × 1024 along with the reference SR-SIM images of the same size are used as input-output pairs for the RED-Net. The network is trained for 100 epochs with the ADAM optimizer and L2 loss.

### 4.5 Architecture and training of U-Net-fairSIM

In U-Net-fairSIM we simply replaced the RED-Net from the RED-fairSIM approach by the U-Net architecture. The U-Net is also trained for 100 epochs with the ADAM optimizer and L2 loss. The complete architecture and parameters of the U-Net are shown in supplementary Fig. 4.

## 5 Hardware and Data availability

During this work we used multiple Nvidia Tesla *P*100 graphics cards. Each network is trained in parallel with 2 Tesla *P*100 to reduce the computational time. The machine learning code is written based on the Python libraries *keras* and *tensorflow*, whereas the *fairSIM* code is written in Java. The code and data which were used during this work can be provided upon request. The generated results are available at https://doi.org/10.5281/zenodo.4095446.

## Acknowledgments

The authors would like to thank Dr. Matthias Fricke and David Pelkmann from the Center for Applied Data Science (CfADS) at Bielefeld University of Applied Sciences for providing access to the GPU compute cluster and Dr. Wolfgang Hubner for providing U2OS cell data with a red fluorescent tubulin stain. We are also thankful to Wadhah Zai el Amri (CfADS) for precursory work on deep learning for the denoising of microscope images. The authors would additionally like to thank Drs. Florian Jug, Carlas Smith, Peter Dedecker and Wim Vandenberg for fruitful discussions on the application of deep learning to SIM reconstruction.

This work was funded by the BMBF (German Federal Ministry of Education and Research) via grant 01IS18041C (consortium project “ITS.ML: Intelligent technical systems via machine learning”). T.H. acknowledges funding by the Deutsche Forschungsgemeinschaft (DFG, German Science Foundation) — project number 415832635. M.M acknowledges funding from the European Union’s Horizon 2020 research and innovation program under the Marie Skłodowska-Curie grant agreements No. 752080.

## Author contributions

Z.H.S. carried out the machine learning work, created the figures, and wrote a large part of the first draft of the manuscript. T.H. and W.S. conceptualized and supervised the research. In addition, they designed the experiments together with Z.H.S. and M.M. The manuscript was refined and extended by T.H., M.M., and W.S. T.-C.W. recorded the raw SIM images and participated in discussions of the denoising approaches. M.M. supported the data reconstruction with fairSIM. P.M.S. supported the machine learning work by running supplementary experiments intended for validation. A.S. and M.S. provided additional ideas from the signal processing perspective. All authors discussed the results and commented on the final manuscript.

## References

[1] Schermelleh, L. et al. Super-resolution microscopy demystified. Nature cell biology 21, 72–84 (2019).

[2] Demmerle, J. et al. Strategic and practical guidelines for successful structured illumination microscopy. Nature Protocols 12, 988 (2017).

[3] Heintzmann, R. & Huser, T. Super-resolution structured illumination microscopy. Chemical Reviews 177, 13890–13908 (2017).

[4] Gustafsson, M. G. Surpassing the lateral resolution limit by a factor of two using structured illumination microscopy. Journal of microscopy 198, 82–87 (2000).

[5] Hirvonen, L. M., Wicker, K., Mandula, O. & Heintzmann, R. Structured illumination microscopy of a living cell. European Biophysics Journal 38, 807–812 (2009).

[6] Kner, P., Chhun, B. B., Griffis, E. R., Winoto, L. & Gustafsson, M. G. Super-resolution video microscopy of live cells by structured illumination. Nature methods 6, 339–342 (2009).

[7] Shao, L., Kner, P., Rego, E. H. & Gustafsson, M. G. Super-resolution 3d microscopy of live whole cells using structured illumination. Nature methods 8, 1044–1046 (2011).

[8] Gao, L. et al. Noninvasive imaging beyond the diffraction limit of 3d dynamics in thickly fluorescent specimens. Cell 151, 1370–1385 (2012).

[9] Fiolka, R., Shao, L., Rego, E. H., Davidson, M. W. & Gustafsson, M. G. Time-lapse two-color 3d imaging of live cells with doubled resolution using structured illumination. Proceedings of the National Academy of Sciences 109, 5311–5315 (2012).

[10] Huang, X. et al. Fast, long-term, super-resolution imaging with hessian structured illumination microscopy. Nature biotechnology 36, 451 (2018).

[11] Markwirth, A. et al. Video-rate multi-color structured illumination microscopy with simultaneous real-time reconstruction. Nature communications 10, 1–11 (2019).

[12] Sandmeyer, A. et al. Dmd-based super-resolution structured illumination microscopy visualizes live cell dynamics at high speed and low cost. bioRxiv 797670 (2019).

[13] Gustafsson, M. G. et al. Three-dimensional resolution doubling in wide-field fluorescence microscopy by structured illumination. Biophysical journal 94, 4957–4970 (2008).

[14] Müller, M., Mönkemöller, V., Hennig, S., Hübner, W. & Huser, T. Open-source image reconstruction of super-resolution structured illumination microscopy data in imagej. Nature communications 7, 1–6 (2016).

[15] Lal, A., Shan, C. & Xi, P. Structured illumination microscopy image reconstruction algorithm. IEEE Journal of Selected Topics in Quantum Electronics 22, 50–63 (2016).

[16] Křížek, P., Lukes, T., Ovesny, M., Fliegel, K. & Hagen, G. M. Simtoolbox: a matlab toolbox for structured illumination fluorescence microscopy. Bioinformatics 32, 318–320 (2016).

[17] Wicker, K., Mandula, O., Best, G., Fiolka, R. & Heintzmann, R. Phase optimisation for structured illumination microscopy. Opt. Express 21, 2032–2049 (ts). URL http://www.opticsexpress.org/abstract.cfm?URI=oe-21-2-2032.

[18] Fan, J., Huang, X., Li, L., Tan, S. & Chen, L. A protocol for structured illumination microscopy with minimal reconstruction artifacts. Biophysics Reports 5, 80–90 (2019).

[19] Hoffman, D. P. & Betzig, E. Tiled reconstruction improves structured illumination microscopy. bioRxiv (2020).

[20] Jin, L. et al. Deep learning enables structured illumination microscopy with low light levels and enhanced speed. Nature Communications 11, 1–7 (2020).

[21] Christensen, C. N., Ward, E. N., Lio, P. & Kaminski, C. F. Ml-sim: A deep neural network for reconstruction of structured illumination microscopy images. arXiv preprint arXiv:2003.11064 (2020).

[22] Ling, C. et al. Fast structured illumination microscopy via deep learning. Photonics Research 8, 1350–1359 (2020).

[23] Weigert, M. et al. Content-aware image restoration: pushing the limits of fluorescence microscopy. Nature methods 15, 1090–1097 (2018).

[24] Sage, D. et al. Quantitative evaluation of software packages for single-molecule localization microscopy. Nature methods 12, 717–724 (2015).

[25] Novak, T., Gajdos, T., Sinkó, J., Szabó, G. & Erdélyi, M. TestSTORM: Versatile simulator software for multimodal super-resolution localization fluorescence microscopy. Scientific Reports 7, 951 (2017). URL http://www.nature.com/articles/s41598-017-01122-7.

[26] Mao, X., Shen, C. & Yang, Y.-B. Image restoration using very deep convolutional encoder-decoder networks with symmetric skip connections. In Advances in neural information processing systems, 2802–2810 (2016).

[27] Lim, B., Son, S., Kim, H., Nah, S. & Mu Lee, K. Enhanced deep residual networks for single image super-resolution. In Proceedings of the IEEE conference on computer vision and pattern recognition workshops, 136–144 (2017).

[28] Zhang, Y. et al. Image super-resolution using very deep residual channel attention networks. In Proceedings of the European Conference on Computer Vision (ECCV), 286–301 (2018).

[29] Hore, A. & Ziou, D. Image quality metrics: Psnr vs. ssim. In 2010 20th international conference on pattern recognition, 2366–2369 (IEEE, 2010).

[30] Ronneberger, O., P. Fischer & Brox, T. U-net: Convolutional networks for biomedical image segmentation. In Medical Image Computing and Computer-Assisted Intervention (MICCAI), vol. 9351 of LNCS, 234–241 (Springer, 2015). URL http://lmb.informatik.uni-freiburg.de/ Publications/2015/RFB15a. (available on arXiv:1505.04597 [cs.CV]).

[31] Abascal, J. F. et al. A residual u-net network with image prior for 3d image denoising. HAL preprint hal-02500664 (2020).

[32] Torrey, L. & Shavlik, J. Transfer learning. In Handbook of research on machine learning applications and trends: algorithms, methods, and techniques, 242–264 (IGI global, 2010).

[33] Howard, J. & Ruder, S. Universal language model fine-tuning for text classification. arXiv preprint arXiv:1801.06146 (2018).

[34] Dabov, K., Foi, A., Katkovnik, V. & Egiazarian, K. Image denoising by sparse 3-d transform-domain collaborative filtering. IEEE Transactions on image processing 16, 2080–2095 (2007).

[35] Lehtinen, J. et al. Noise2Noise: Learning image restoration without clean data. In Dy, J. & Krause, A. (eds.) Proceedings of the 35th International Conference on Machine Learning, vol. 80 of Proceedings of Machine Learning Research, 2965–2974 (PMLR, Stockholmsmassan, Stockholm Sweden, 2018).

[36] Zhu, J.-Y., Park, T., Isola, P. & Efros, A. A. Unpaired image-to-image translation using cycle-consistent adversarial networks. In Proceedings of the IEEE international conference on computer vision, 2223–2232 (2017).

[37] Karras, C. et al. Successful optimization of reconstruction parameters in structured illumination microscopy-a practical guide. Optics Communications 436, 69–75 (2019).

