## Supplementary material for "Deep-learning based denoising and reconstruction of super-resolution structured illumination microscopy images"

### SR-REDSIM Architecture

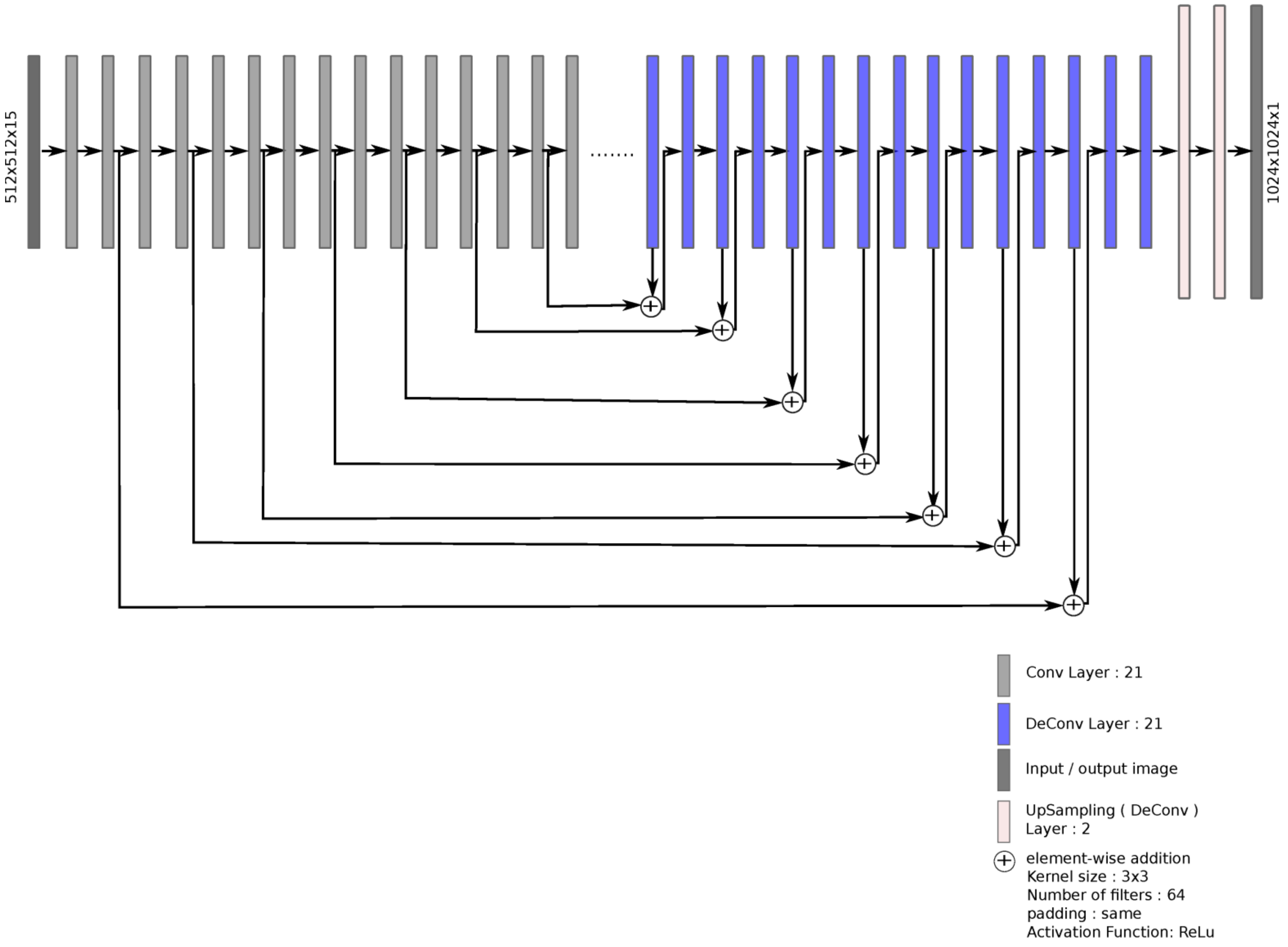


**Supplementary figure 1:** This figure shows the architecture of SR-REDSIM. The network is composed of three different blocks. The encoding and decoding block contain 21 convolutional and deconvolutional layers, respectively, whereas the up-sampling blocks consist only of two up-sampling layers. This architecture was used in the SR-REDSIM method to denoise and reconstruct the raw SIM images.

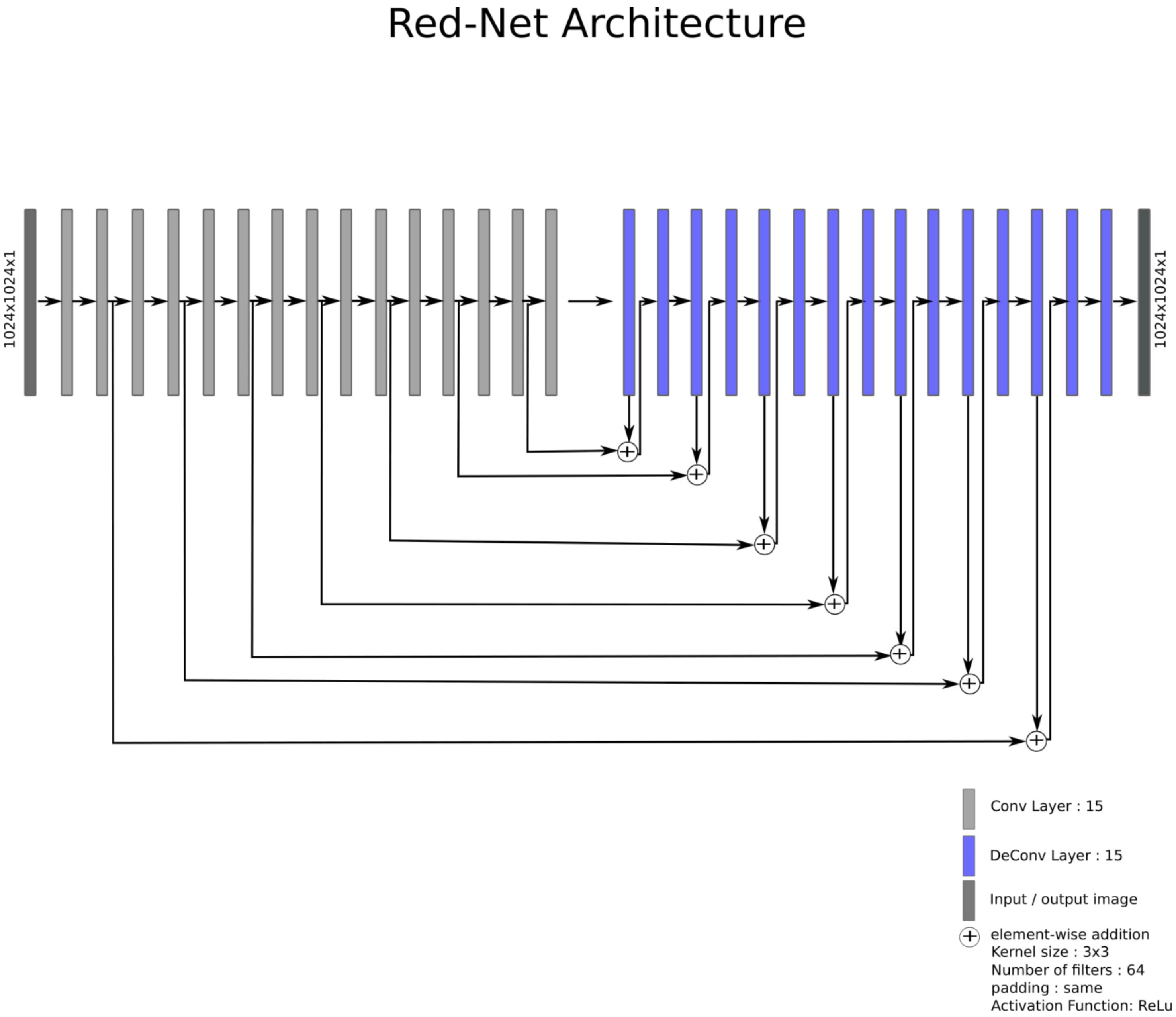


**Supplementary figure 2:** The complete architecture of the “Residual Encoder-Decoder Network” is shown in this figure. The architecture contains 15 convolution and 15 deconvolutional layers along with the additive symmetric skip connection layers. This architecture was used in the RED-fairSIM and preRED-fairSIM methods for denoising.

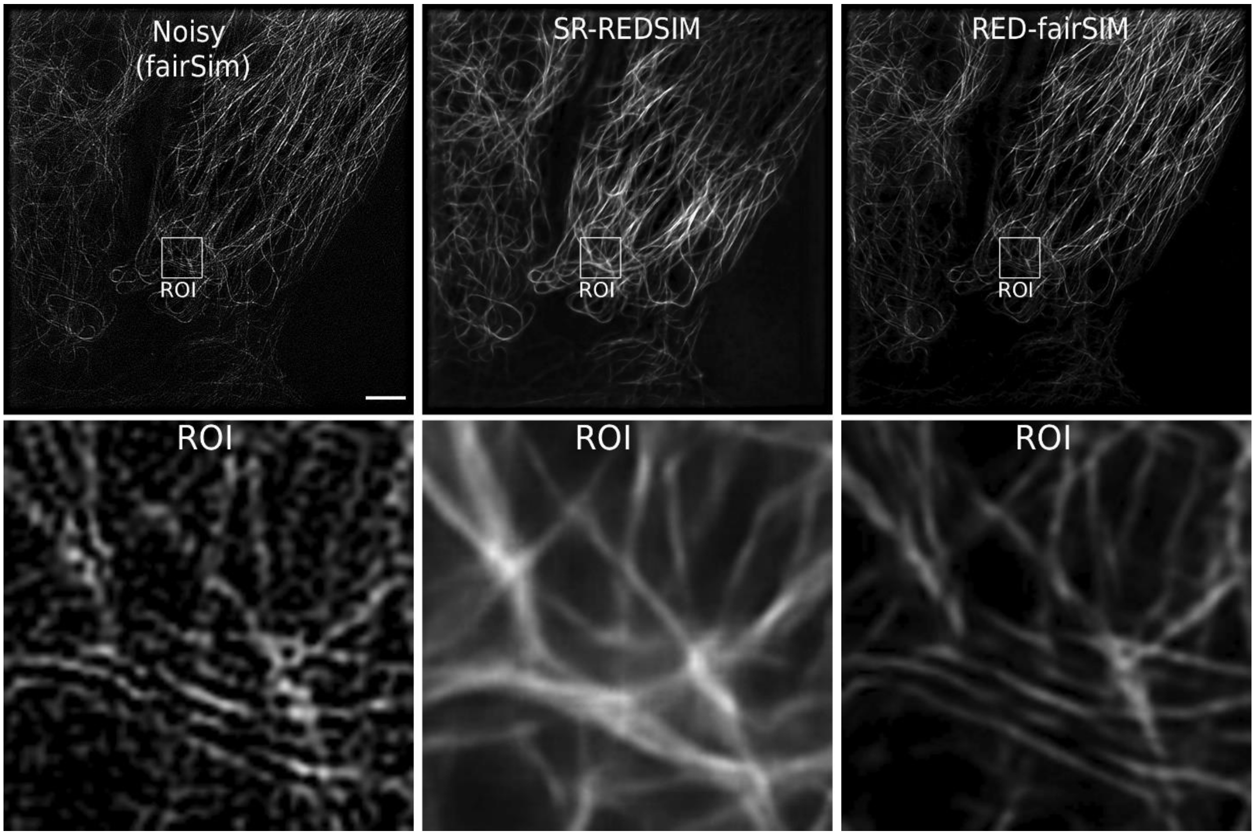


**Supplementary figure 3:** These SR-SIM images show the difference between the output of the SR-REDSIM and RED-fairSIM methods when applied to imaging conditions that the underlying network was not trained for. To evaluate the generalization capabilities of these methods, we again collected tubulin structure (on U2OS cells), but with a different excitation wavelength. Here, the cell is illuminated by light with a wavelength of 642 nm instead of 488 nm (the latter used for the images in the training set). The different wavelength also changes the spatial frequency of the SIM patterns. This cell structure with unseen illumination properties is then propagated through the pretrained models of both SR-REDSIM and RED-fairSIM. The resulting SR-SIM image shows that RED-fairSIM is more robust against changed microscope settings than SR-REDSIM. Scale bar: 4 µm.

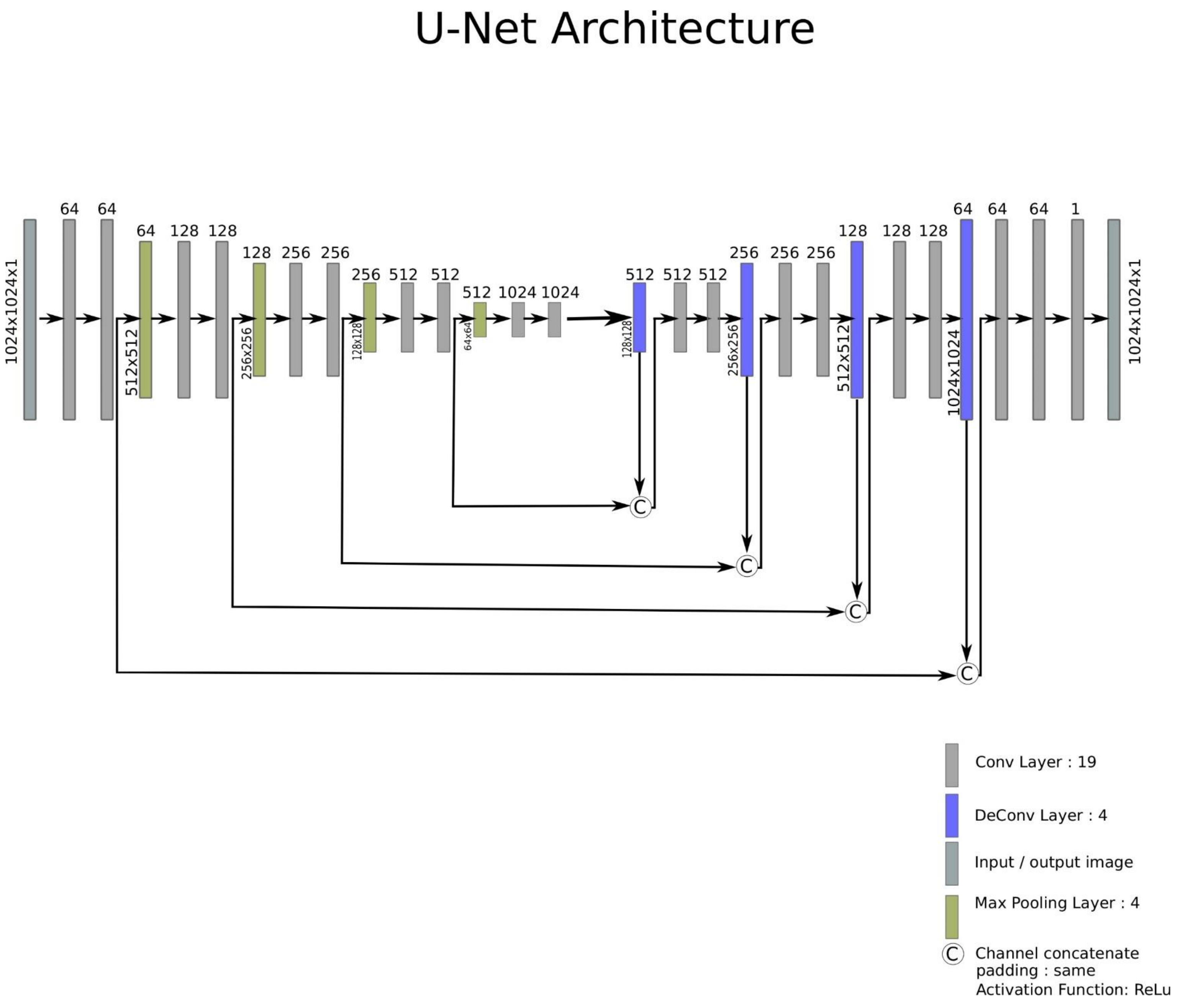


**Supplementary figure 4:** This figure illustrates the architecture of U-Net. This architecture was used in the U-Net-fairSIM pipeline. In U-Net-fairSIM, U-Net is combined with the fairSIM algorithm to denoise noisy SR-SIM images.

| Mean PSNR and SSIM values for noise level 4 | | | | |
|---|---|---|---|---|
| | PSNR | SSIM | PSNR with Augmentation Approach | SSIM with Augmentation Approach |
| SR-REDSIM | 26.62 | 0.69 | 26.37 | 0.66 |
| U-Net-fairSIM | 26.80 | 0.68 | 28.05 | 0.71 |
| RED-fairSIM | 27.97 | 0.71 | 28.09 | 0.72 |

**Supplementary table 1:** This table reports the average PSNR and SSIM values (on the test data) of several models from different methods with and without image augmentation in the training data. In image augmentation, each image was used twice for training by adding a version rotated by an angle of 180° to the training set. This approach doubles the amount of training data from 2025 to 4050 images. The test data stays unchanged and is not augmented. The augmentation approach shows (slight) improvements in the PSNR and SSIM results of U-Net-fairSIM and RED-fairSIM, however, on the other side it doubles the required training time.

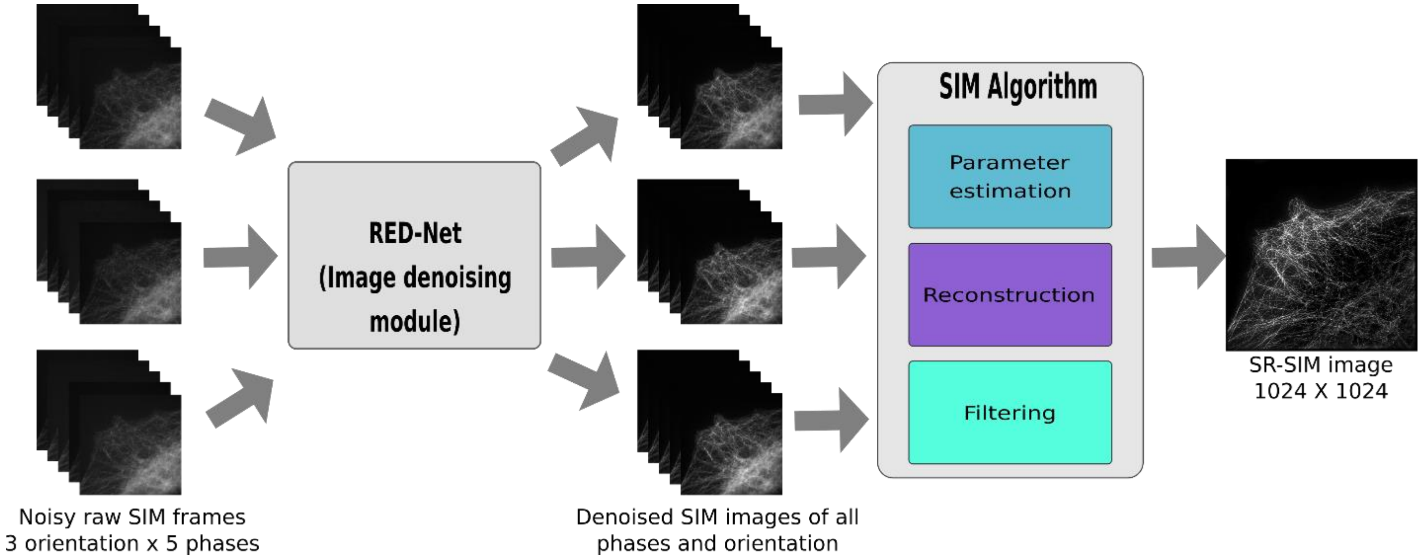


**Supplementary figure 5:** This figure shows the complete pipeline of the preRED-fairSIM method. In this pipeline, the raw SIM images (512x512, (width x height)) of all phases and orientations are denoised separately with the RED-Net architecture. The complete architecture of RED-Net is shown in supplementary figure 2. The denoised SIM images of each phase and orientation are then propagated into the fairSIM software in the form of a stack (15 frames) to reconstruct the super-resolution SIM image.

| Mean PSNR and SSIM values | | | | | | |
|---|---|---|---|---|---|---|
| **Dataset** | **RED-30** | | **Transfer learning** | | **Fine-tuning** | |
| | PSNR | SSIM | PSNR | SSIM | PSNR | SSIM |
| **Phase 0 Orientation 0** | 33.22 | 0.90 | - | - | | |
| **Phase 0 Orientation 1** | - | | 33.43 | 0.90 | | |
| **Phase 0 Orientation 2** | - | | 33.15 | 0.90 | | |
| **Phase 1 Orientation 0** | - | | 33.01 | 0.89 | 33.12 | 0.89 |
| **Phase 1 Orientation 1** | - | | 33.20 | 0.90 | 33.58 | 0.90 |
| **Phase 1 Orientation 2** | - | | 33.32 | 0.90 | 33.28 | 0.89 |
| **Phase 2 Orientation 0** | - | | 33.17 | 0.90 | 33.60 | 0.90 |
| **Phase 2 Orientation 1** | - | | 33.29 | 0.90 | 33.12 | 0.89 |
| **Phase 2 Orientation 2** | - | | 32.99 | 0.88 | 33.36 | 0.90 |
| **Phase 3 Orientation 0** | - | | 33.33 | 0.89 | 33.53 | 0.90 |
| **Phase 3 Orientation 1** | - | | 33.40 | 0.90 | 33.44 | 0.90 |
| **Phase 3 Orientation 2** | - | | 33.19 | 0.89 | 33.16 | 0.89 |
| **Phase 4 Orientation 0** | - | | 33.07 | 0.88 | 33.27 | 0.89 |
| **Phase 4 Orientation 1** | - | | 33.59 | 0.90 | 33.61 | 0.90 |
| **Phase 4 Orientation 2** | - | | 33.41 | 0.90 | 33.01 | 0.89 |

**Supplementary table 2:** This table shows the mean PSNR and SSIM values of the preRED-fairSIM method on the test data. The 30-layer RED-Net is first trained on phase 0 and orientation 0 and afterwards tested on the data from different phases and orientations. During the transfer learning, the trained model shows on the data of other phases and orientations similar PSNR and SSIM values (third column) as on phase 0/orientation 0 (second column). Furthermore, in a subsequent experiment, the pretrained network was fine-tuned by retraining the first five convolutional layers and the last five deconvolutional layers for 20 epochs. The fine-tuning step did not improve the results in any significant way (fourth column).

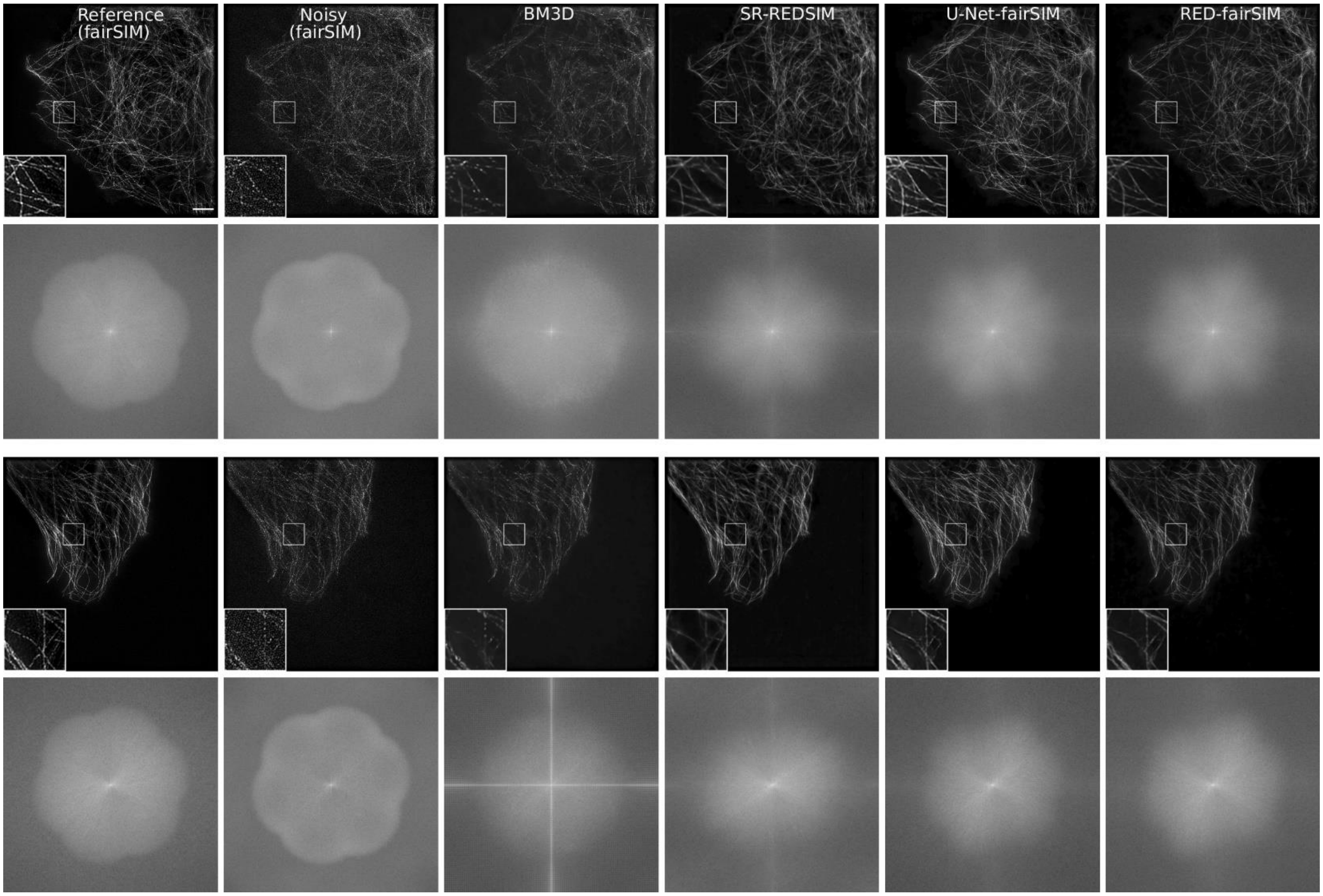


**Supplementary figure 6:** This figure shows the reconstructed SR-SIM images of two test samples with different methods. The Fourier spectrum of each SR-SIM image is shown directly below. Each image contains an enlarged region of interest (ROI) at the lower-left bottom. The analysis of ROI of all the methods clearly shows that the results of RED-fairSIM (sixth column) are smoother and more faithful compared to all other methods. Similarly, the Fourier spectra of the RED-fairSIM and U-Net-fairSIM (fifth column) results do not show any additional artifacts in the Fourier space. The SR-SIM images and ROI of SR-REDSIM (fourth column) also show good results however, there are some artifacts in the high-frequency region of the Fourier spectrum. The ROI of BM3D (third column) shows a suppressed cell structure in both of the resultant images. Furthermore, the Fourier spectrum of the BM3D result for the second test sample shows artifacts in both low and high-frequency regions. Scale bar: 4 µm.
